# Neural Correlates of Self-Referential Belief Processes

**DOI:** 10.1101/2023.10.05.561046

**Authors:** Emily Bruns, Immanuel Scholz, Georgia Koppe, Peter Kirsch, Martin Fungisai Gerchen

## Abstract

**Background:** Belief processing as well as self-referential processing have both been consistently associated with cortical midline structures. In addition, seminal neuroimaging papers have implicated cortical regions such as the vmPFC in *general* belief processing. However, the neural correlates of *self-referential* belief are yet to be investigated in functional magnetic resonance imaging (fMRI).

**Methods:** In this fMRI study, we presented 120 statements with trait adjectives as target words to N=27 young healthy participants and asked them to judge whether they believed that these trait adjectives applied to themselves, a self-chosen close person, or a public person (the German chancellor at that time). Participants subsequently rated how certain (0-100%) they were in their judgment.

**Results:** As expected, self-referential processing evoked a large cluster in the vmPFC, ACC and dmPFC. For belief, we found an activated cluster in the vmPFC during statement presentation, which partly overlapped with the cluster for self-referential processing. The cluster for self-belief vs. disbelief was similar in location and size to the cluster for general belief processing and distinct from the cluster for self-referential processing. We also found dmPFC activation for uncertainty in belief evaluations.

**Discussion:** We successfully replicated vmPFC involvement in belief processing and found a common neural correlate for belief and self-belief in the vmPFC. The activation clusters for self-belief versus self-referential processing were distinct, implying distinct neural processes. This insight will prove relevant for investigations in clinical populations with aberrant (self-)belief processing. Furthermore, we replicated the role of the dmPFC in uncertainty, supporting a dual neural process model of belief and certainty.

## Introduction

Belief is a substantial aspect of everyday human experience and crucial in shaping subjective reality, emotion, and behavior (Harris et al., 2008; James, 1890/1910). It has far-reaching implications for the public and political domain, including societal polarization and the spread of conspiracy theories (Bowen et al., 2023; Gerchen et al., 2023; Haghtalab et al., 2021), to inflexible individual convictions in mental disorders like delusion and depression.

While belief has not been a dominant focus of psychological research in the last decades, it is a traditional psychological construct. William James conceptualized belief as “the mental state or function of cognizing reality” and “a sort of feeling more allied to the emotions than to anything else” (1890/1910; p. 283), highlighting its emotive and subjective component. The nature of belief processes has been intensively discussed in the early psychological literature. Already in 1859, Alexander Bain postulated that the opposite of belief is doubt or uncertainty: “The real opposite of belief as a state of mind is not disbelief, but doubt, uncertainty; and the close alliance between this and the emotion of fear is stamped on every language.” (Bain, 1859, p. 574), introducing the dimension of certainty and highlighting its importance for an individual’s emotional state. The more recent psychological literature particularly focuses on the relationship of belief and disbelief processes (Asp 2012; Asp, 2022; Gilbert, 1991; Gilbert, 1993; Hasson, 2005).

Nowadays, neuroimaging approaches are one of the most promising ways to further elucidate the processes underlying belief (see for example Harris et al., 2008; Harris et al., 2009; Marques et al., 2009; Douglas et al., 2013; Seitz, 2022; Seitz & Angel, 2012; Seitz et al., 2018). In a seminal fMRI study by Harris et al. (2008), participants were shown statements from different categories (autobiographical, mathematical, geographical, religious, ethical, semantic, and factual), and judged whether they were “true”, “false”, or “undecidable”. In this study, the authors associated the ventromedial prefrontal cortex (vmPFC) with belief. The vmPFC has been linked to behavioral modulation to rewards and goals (Grabenhorst & Rolls, 2011; Hornak et al., 2004; Matsumoto & Tanaka 2004), emotional saliency (Goel & Dolan, 2003) as well as the recruitment of somatic markers in decision-making (Bechara et al., 2000; Damasio, 1996). Notably, in the context of belief processing, the vmPFC has also been proposed as a neural correlate of a “false tagging” process, phenomenally manifesting as doubt and skepticism (Asp et al., 2012). Interestingly, this notion is consistent with a larger deflection of vmPFC activity for disbelief than for belief, which creates the positive contrast effect for belief (Harris et al., 2008; Gerchen et al., 2023; see also Supplement, Figure 1). For disbelief, Harris et al. (2008) found effects in the left inferior frontal gyrus, anterior insula, dorsal anterior cingulate (dACC), superior frontal gyrus, and superior parietal lobule. An important limitation of the study by Harris et al is, however, that belief was counfounded with (un)certainty, since the authors did not assess certainty in “true” and “false” belief evaluations, or uncertainty in “undecidable” statements.

**Figure 1.**
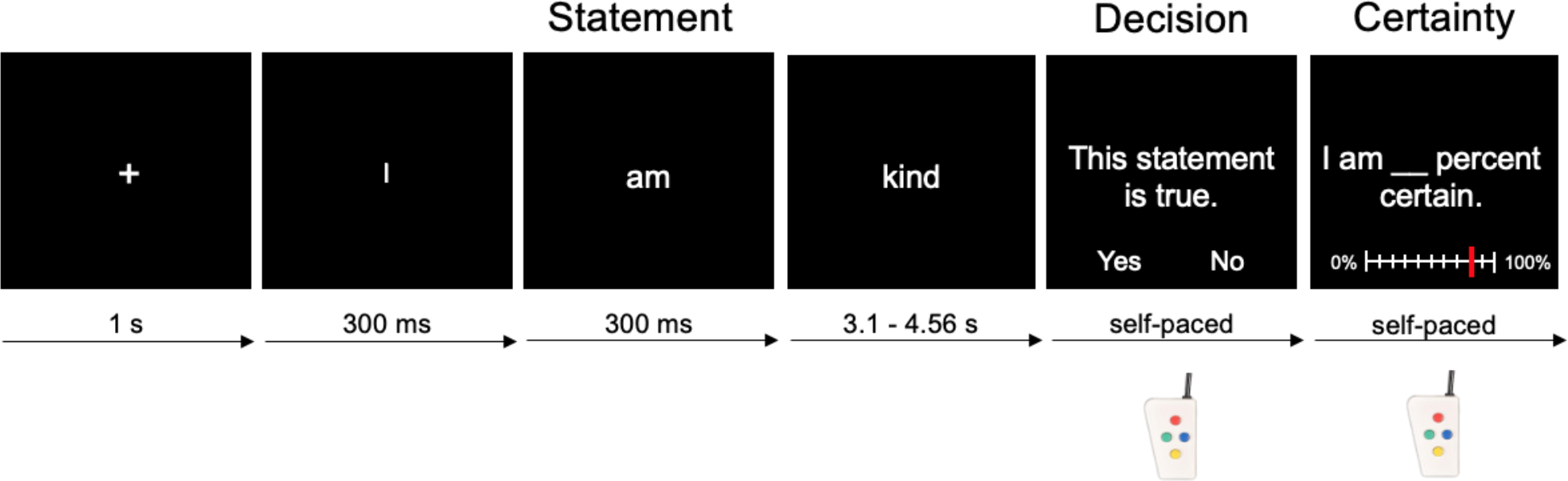
Statement Judgment Task. 120 statements containing trait adjectives as target words were presented in random order. The target word was presented for a jittered duration of 3.1-4.56 s. Participants were not required to react during the rapid visual presentation but were subsequently prompted to make a forced-choice, self-paced binary decision about the truth / falsehood of that statement. Thereafter, participants evaluated their certainty in the previous belief evaluation on a visual analog scale from 0-100% in their own time. The original statements were presented in German.

Based on Harris et al.’s (2008) paradigm, Gerchen et al. (2023) conducted an fMRI study to investigate general belief in factual, political, religious, conspiracy, and superstitious, but not in self-referential, statements. In contrast to Harris et al. (2008), we (Gerchen et al. 2023) temporally separated statement presentation from truth evaluation and applied a forced choice decision “true”/ “false” decision with a separate certainty assessment. Interestingly, we found similar vmPFC activation for belief as Harris et al. (2008), but at the time of judgment and not during the preceding statement presentation. During statement presentation, we identified activity in bilateral dorsolateral prefrontal cortex (dlPFC), left superior parietal cortex, and left lateral frontopolar cortex (left lFPC) for statements evaluated as “true”. For disbelief, we found activation in the temporal pole (TP), a structure serving as an amodal semantic hub (Chen et al., 2017; Patterson et al., 2007). The TP is also involved in the processing of semantic meaning (Tsapkini et al., 2011; Vandenberghe et al., 2002) and semantic memory (Bonner & Price, 2013; Simmons & Martin, 2009), and has been implicated in belief-laden as opposed to belief-neutral reasoning (Goel & Dolan, 2003). Notably, the strongest effect we found was activation in the dorsomedial prefrontal cortex (dmPFC) associated with uncertainty, which we interpreted as a marker of the “subjective epistemic risk” of a statement. Based on this finding, we propose dual neural belief processes with veracity/falsehood and uncertainty/certainty as two distinguishable factors.

An important further question is whether belief processes are general processes or whether different functions are involved in belief in different domains. For example, comparing religious and non-religious belief, Harris et al. (2009) found that religious belief had higher associations with brain processes involved in emotion, self-representation, and cognitive conflict. In another study, Howlett & Paulus (2015) found that testability of statements was associated with activation in the dlPFC and posterior cingulate cortex.

One of the most relevant potential distinctions of belief domains in humans is the differentiation between self-referential and non-self-referential belief. Better understanding of self-referential belief processing is especially important for laying the foundation for future studies in clinical contexts, as self-beliefs and aberrant self-referential processing are highly relevant in mental disorders. In fact, negative self-beliefs are associated with mental health issues across disorders (e.g., depression, psychosis, eating disorders) and cultures and ethnic groups (Cooper & Cowen, 2009; Dozois & Rnic, 2015; Nguyen et al., 2019; Orth & Robbins, 2013; Smith et al., 2006). For example, self-referential cognition is a pivotal component of Beck’s cognitive triad of depression (1967), and aberrant beliefs and belief updating are hallmarks of psychosis (Fletcher & Frith, 2009; Kube & Rosenkrantz, 2021). It is thus imperative that future clinical studies on the diagnosis and treatment (for psychosis see Pott & Schillbach, 2022) of these disorders consider how changes in belief processing are represented at the neural level.

For the first study aiming to make this differentiation, Han et al. (2017) developed an fMRI paradigm based on trait adjectives describing the participant or a gender-matched celebrity. However, they did not focus on the differentiation between self-referential and non-self-referential belief, but rather on the contrast of “believing” vs. “thinking”. For this contrast, they found stronger activation in the left anterior insula (AI) / inferior frontal cortex (IFC) as well as greater intrinsic connectivity between the left AI / IFC and the left occipital cortex. Regarding self-belief, they found that, in the belief condition, self-judgments were associated with greater activation in the left AI / IFC as well as greater functional connectivity between the mPFC and the left occipital cortex than valence-judgments during judgment. Using the same paradigm, Gao et al. (2022) compared neural belief processes between Chinese and Danish participants. For both cultural groups, they identified the mPFC in association with general belief over both conditions. In an analysis secondary to their main research question, they found that self-vs. celebrity-related belief was associated with activation in the vmPFC.

The finding by Gao et al. (2022) of a stronger self-referential belief effect in the vmPFC emphasizes the question whether self-referential and non-self-referential belief processes are based on similar or distinct neural processes. While belief processing as well as self-referential processing (Kim, 2012; Kurczek et al., 2015; Lemogne et al., 2011; Northoff et al., 2006) have both been associated with cortical midline structures (CMS) overlapping in the vmPFC, further and more explicit investigation is warranted as to how this rough correspondence is reflected in actual activations, and as to whether the involved CMS differ for belief of self-referential content.

In the present study, our aim was therefore to investigate the neural correlates of self-referential belief in comparison to non-self-referential belief. To address this question, we designed a paradigm related to Han et al. (2017). In our paradigm, participants judged short statements about themselves, a close person (a self-chosen close friend or family member), and a public person (the long-time German chancellor at the time of the experiment, Angela Merkel). As in our former work (Gerchen et al., 2023), responses were given as a binary forced-choice belief decision followed by a certainty rating for the decision (Figure 1). This approach also allowed us to compare the obtained results for self-referential belief to those of Gerchen et al. (2023) for general belief and investigate differences as well as the replication of effects.

## Materials & Methods

### Participants

In total, N=32 young healthy participants were recruited for the study. The participants were mainly university students from the German cities Mannheim, Heidelberg, and Freiburg. All participants were screened by telephone for exclusion criteria prior to participation. Exclusion criteria were left-handedness, uncorrected impaired vision, a diagnosis of a neurological or mental disorder, pregnancy, implants or metal parts in the body, and claustrophobia. The data sets of five participants had to be excluded due to anatomical incidental findings (1), inadequate language skills (1), and excessive head motion (3). Thus, N=27 participants (14 women; average age 23.3 years; age range 19-38 years) were included in the analyses. All participants were fully informed about the study and provided written informed consent. After the experiment, participants received either a reimbursement of 30€ or partial course credit at their university. The study was approved by the ethics committee of the Medical Faculty Mannheim, Heidelberg University, Germany (2020-649N) and procedures complied with the Declaration of Helsinki of the World Medical Association.

### Data Acquisition

MRI measurements were conducted on a Siemens Magnetom TrioTIM 3T scanner (Siemens Healthineers, Erlangen, Germany) at the Central Institute of Mental Health in Mannheim, Germany. Data for each participant was acquired in a single session that consisted of an anatomical measurement, a 6-minute resting state scan, and the task. Anatomical images were acquired with a magnetization prepared rapid gradient echo (MPRAGE) sequence with repetition time TR=2.3s, echo time TE=3.03ms, flip angle=9° and an isotropic resolution of 1×1×1mm. Functional images were acquired with an Echo Planar Imaging (EPI) sequence with TR=1.64s, TE=30ms, flip angle=73° and GRAPPA factor 2 in 30 slices of 3mm thickness with 1mm gap, an in-plane resolution of 3×3mm, and field of view FoV=192mm. In addition, respiratory and cardiac activity was recorded during functional measurements with built-in equipment.

### Experimental Task

The experiment consisted of three different conditions: belief about the self, a self-chosen close friend or family member (“close person”), and the previous German chancellor, Angela Merkel (“public person”). Two other-conditions were used to prevent a mono-operation bias (Cole et al., 1981) and to assess potential effects of intimacy and autobiographical processing. Angela Merkel was chosen because she is a well-known public figure, whom we did not expect the participants to know personally (which none of the participants did, according to their self-report). All three conditions aimed to trigger subjective belief processing of personality traits (e.g., “I am / [Name of close person] is / Merkel is lazy.”). The experiment used a rapid serial visual presentation paradigm. First, a fixation cross appeared on a monitor for 1 s, followed by three individual words each shown for 300 ms. The final target word, which conveyed the meaning of the statement, was presented for a jittered period of 3.1 to 4.56 s. Subsequently, the participants were prompted to a self-paced belief evaluation of the statement (“This statement is true.”; German: “Diese Aussage ist wahr.”). This prompt was included, as Wiswede et al. (2013) postulated truth evaluation to depend on task demands and an evaluative mindset rather than being an automatic process. The “yes” or “no” answers were randomly assigned to the left or right button of a small hand-held four-button controller to prevent confounding preparatory motor activation during the evaluation period. Afterwards, participants rated how certain they were in their evaluation (“I am … percent certain.”; German: “Ich bin mir sicher zu …”) on a visual analogue scale from 0% to 100% in their own time. The start value was set to 50% and participants could navigate to their certainty value of choice using the hand-held controller. An exemplary trial is shown in Figure 1. Participants completed 120 trials in the fMRI scanner. The average duration of the task was 26:13 ± 2:52 min (range: 23:14-37:01 min).

A pool of trait adjectives was generated specifically for this paradigm because there were no preexisting trait-adjective lists normed for endorsement of the trait adjective for the self, a close person, and a public figure. We aimed for an even distribution of statements rated as true and false, and thus included negative, neutral, and positive trait adjectives. The final 80 items drew on a pool of trait adjectives from Lazzari et al. (1978) and King (1983), of likability-rated items from Dumas et al. (2002), Schonbach (1972) and Alicke (1985), and on self-generated trait adjectives.

We designed our experiment to be conducted with two parallel statement sets subsequently with magnetoencephalography (MEG) and fMRI. To split the statements into two sets, 9 participants were recruited to pre-test the paradigm including all 80 target words. Based on the pre-test data, the pool of target words was partitioned into two maximally equivalent and internally heterogeneous sets of 40 items for the parallel test versions using Papenberg and Klau’s (2021) anti-clustering method. The parallel versions were used for MEG and fMRI and counterbalanced over participants. The order of items within a set at presentation was random. In this paper, only the fMRI data is presented.

### Data Analysis

Data analyses were conducted in MATLAB (R2017a; The MathWorks, Inc., Natick, Massachusetts, USA). fMRI data was analyzed with SPM12 (v7219; Wellcome Center for Human Neuroimaging, London, United Kingdom).

### fMRI Preprocessing

The anatomical images were segmented and normalized to the SPM12 TPM MNI template. Functional images were slice-time corrected, realigned to the mean image, co-registered to the anatomical image and normalized to MNI space by applying the forward deformation matrix estimated from the anatomical image. In this step, the functional data was rescaled to 3×3×3mm isotropic resolution and subsequently smoothed with a three-dimensional Gaussian kernel with full width at half maximum FWHM=8×8×8mm. In addition, nuisance signals from regions containing white matter (WM) and cerebrospinal fluid (CSF) were estimated, the ART toolbox (http://www.nitrc.org/projects/artifact_detect) was used to identify volumes affected by movements (framewise displacement FD>0.5mm; scan-to-scan global signal change z>4), and the TAPAS PhysIO toolbox (Kasper et al., 2017) was used to estimate physiological nuisance regressors based on the acquired cardiac and respiratory signals.

### fMRI First Level Analyses

fMRI first level analyses were conducted using general linear models (GLMs). The main first level model to estimate self-referential and belief processes contained 6 experimental regressors of statement presentations (belief and disbelief for the self, close person, and public person conditions, respectively) convolved with the canonical hemodynamic response function (cHRF). Based on this model, the contrasts ‘self vs. other’ (other including the close and public persons) over belief and disbelief statements, and ‘belief vs. disbelief’ over all persons as well as within the conditions were estimated. The first level model for estimating uncertainty effects contained three experimental regressors for statement presentations in the conditions (self, close person, public person) convolved with the cHRF as well as their parametric modulation by trial-wise certainty ratings. Uncertainty effects were estimated by negative contrasts on the parametric modulation regressors over all conditions as well as within the conditions.

Both first-level design matrices contained regressors of the time courses of fixation cross presentations and button presses, and the decision and certainty phase convolved with the cHRF, as well as six motion regressors, WM and CSF regressors, dummy regressors for movement-affected time points, and the physiological nuisance regressors.

### fMRI Second Level Analyses

Participant-specific contrast maps from the first level analyses were entered into second level analyses with one-sample t-tests. All second level models contained sex as covariate. A significance threshold of p=0.05 cluster-level FWE-corrected with cluster-defining threshold (CDT) of p=0.001 uncorrected was applied in all second level analyses.

### Overlap with Gerchen et al. (2023)

For descriptive replication analyses, we plotted the overlap of the identified activations with the ‘belief > disbelief’ effect in the vmPFC and the uncertainty effect in the dmPFC we previously identified in a separate sample with belief judgements on statements about facts, politics, religion, conspiracy theories, and superstition (Gerchen et al., 2023). Here, we applied to the data of Gerchen et al. (2023) the original reported threshold for the vmPFC effect (p=0.05 cluster-level FWE significant with CDT=0.001 unc.) and a stricter than originally reported significance threshold for the dmPFC effect (p=0.05 whole-brain FWE corr.) for illustrative reasons.

## Results

### Behavior

On average, 60.56 ± 7.11% (inter-subject range 44-72%) of statements were judged as true. The self-referential condition showed the highest (65.3 ± 9.56%) and the public-person condition the lowest (53.47 ± 7.64%) rate of belief (paired-sample t-test t(26) = 6.58, p<0.001, g=1.23). 62.89 ± 9.08% of statements about the close person were believed to be true, revealing a significant difference only to the public person (comparison paired-sample t-test t(26) = -4.93, p<0.001, g=-0.92) but not the self (paired-sample t-test t(26) = 1.6, p=0.1221, g=0.3), Participants were 72.99 ± 8.07% (range 56-86%) certain in their belief evaluation on average. They were most certain judging their chosen close person (79.36 ± 8.85%) and least certain judging the public person (62.14 ± 14.68%; comparison paired-sample t-test t(26) = 5.91, p<0.001, g=1.10). On average, participants were 77.4% ± 8.51% certain in belief evaluations about themselves, which was significantly higher than certainty in evaluations about the public (t(26)=4.93, p<0.001, g=0.92 but not the close person (t(26) = -2.01, p = 0.0544, g = -0.38)). Overall, certainty was significantly higher in statements rated as true (75.09 ± 8.47%) than in those believed to be false (69.83 ± 8.42%; paired-sample t-test t(26) = 5.59, p<0.001, g=1.04).

When we compared the two item sets, they did not differ in the rate of belief in the self-referential and close person conditions. However, the rate of belief differed in the public person condition. For certainty, there were no differences between the sets in any of the three conditions (see Supplementary Table 3).

### fMRI self vs. other

Testing the contrast ‘self > other’ (other including the close and public persons) resulted in the largest cluster of activation found in this experiment. Namely, we identified a significant cluster centered in the vmPFC and ACC, which further reached into the dmPFC, corpus callosum and subcortical structures in the diencephalon (Figure 2).

**Figure 2.**
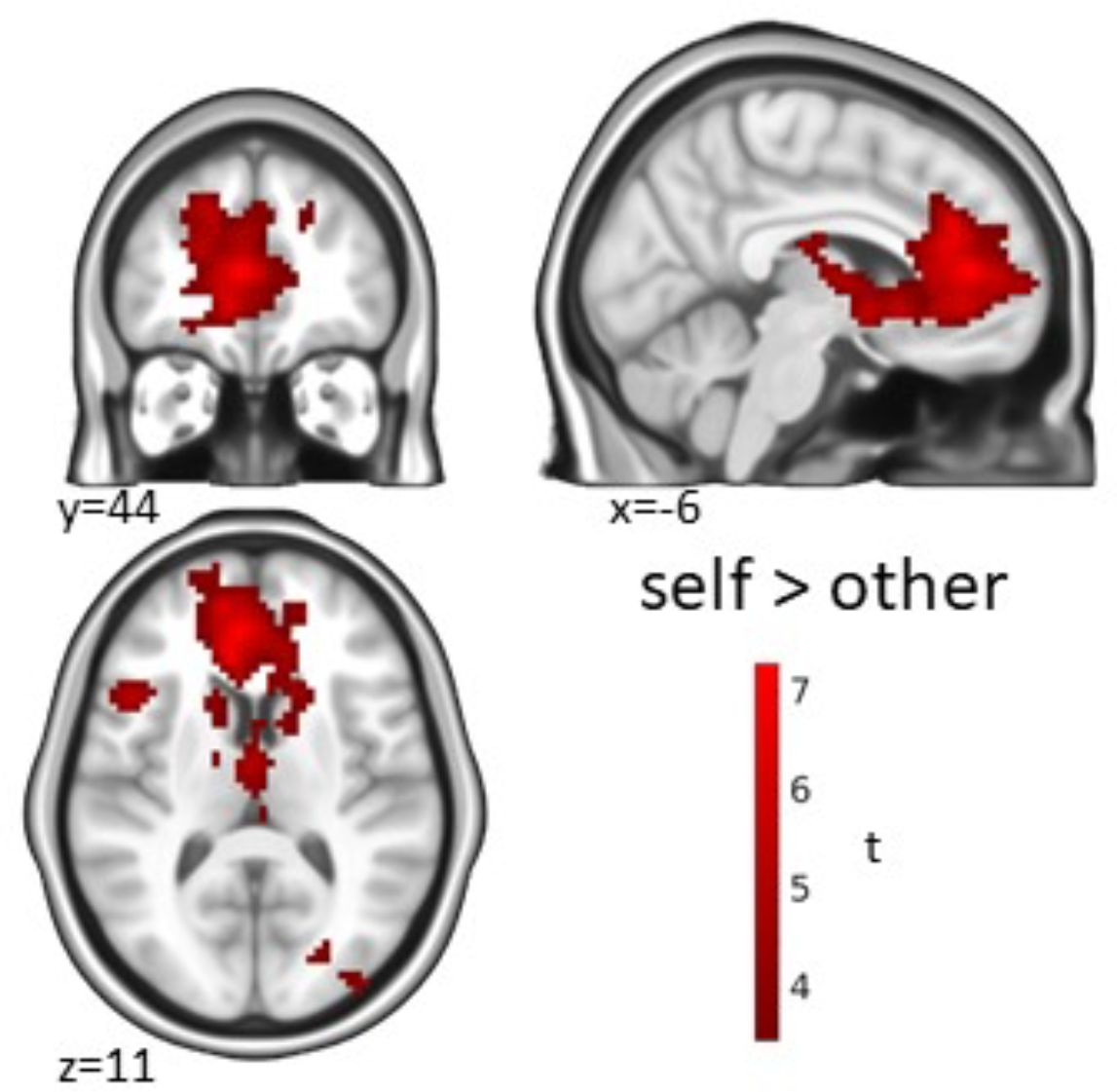
Self-referential processing. Activation for the contrast ‘self > other’ in the statement phase. Significance threshold p=0.05 cluster-level FWE-corrected with a cluster-defining threshold (CDT) of p=0.001 unc.

### fMRI belief vs. disbelief

For the contrast ‘belief > disbelief’, we found a significant cluster in the ACC and vmPFC across all conditions (Figure 3a). As in Gerchen et al. (2023), this ‘belief’ effect was based on stronger deactivation of the vmPFC cluster in the disbelief condition compared to the belief condition (Supplementary, Figure 1), but was present during the presentation phase of the experiment. While we did not find significant differences for the contrast ‘belief > disbelief’ between conditions, the within-condition contrast ‘belief > disbelief’ was only significant in the self-referential condition but not for the close or public persons. The cluster identified in this contrast (‘self belief > self disbelief’) involves the ACC, vmPFC and a small part of the genu of the corpus callosum and is very similar to the cluster for general ‘belief > disbelief’ in shape and localization (Figure 3b). Moreover, it partly overlapped at the edges with the activation for the contrast ‘self > other’ shown in Figure 2 but was distinct from it in size and extent (see Supplementary Figure 3). Exploratively, small subcortical clusters were found in the amygdala and ventral striatum (n.s., cluster-level p=0.09; see Table 2). The activation for self-referential belief showed substantial overlap with the findings on general belief from the fMRI study by Gerchen et al. (2023) and involved the same cortical regions (ACC, vmPFC; see Figure 5a). The contrast ‘disbelief > belief’ revealed no significant clusters.

**Table 1.** Self-referential processing. fMRI results for the contrast ‘self > other’ in the statement phase. Significance threshold p=0.05 cluster-level FWE-corrected with a cluster-defining threshold (CDT) of p=0.001 unc.

| cluster-level |  | peak-level |  |  |
| --- | --- | --- | --- | --- |
| p<sub>FWE</sub> | k<sub>E</sub> | p<sub>FWE</sub> | t | MNI (mm; x, y, z) |
| **self > other** |  |  |  |  |
| <0.001 | 3523 | <0.001 | 10 | -6, 44, 11 |
|  |  | <0.001 | 8.30 | -6, 32, -4 |
|  |  | <0.001 | 8.28 | -9, 5, -7 |
| <0.001 | 643 | 0.03 | 6.23 | 42, -55, -43 |
|  |  | 0.065 | 5.85 | 24, -73, -22 |
|  |  | 0.067 | 5.83 | 45, -64, -34 |
| <0.001 | 305 | 0.25 | 5.14 | -27, -70, -25 |
|  |  | 0.29 | 5.05 | -45, -67, -31 |
|  |  | 0.73 | 4.38 | -45, -58, -43 |

**Table 2.** Belief processing. fMRI results for the contrast ‘belief > disbelief’ over all conditions and the contrast ‘self belief > self disbelief’ in the statement phase. In this case an explorative significance threshold of p=0.10 cluster-level FWE-corrected with a cluster-defining threshold (CDT) of p=0.001 unc. was used to include the small subcortical clusters in the table. Explorative results with 0.05 > p > 0.10 are shown in gray.

| cluster-level |  | peak-level |  |  |
| --- | --- | --- | --- | --- |
| p<sub>FWE</sub> | k<sub>E</sub> | p<sub>FWE</sub> | t | MNI<br>(mm; x, y, z) |
| **belief > disbelief** |  |  |  |  |
| <0.001 | 224 | 0.44 | 4.96 | -6, 38, -10 |
|  |  | 0.59 | 4.75 | -12, 32, -16 |
|  |  | 0.69 | 4.62 | -9, 53, -4 |
| **self belief > self**<br>**disbelief** |  |  |  |  |
| 0.096 | 60 | 0.21 | 5.36 | -18, -10, -19 |
|  |  | 0.94 | 4.12 | -12, -1, -13 |
| <0.001 | 269 | 0.21 | 5.36 | -6, 29, -16 |
|  |  | 0.32 | 5.13 | 6, 38, -16 |
|  |  | 0.38 | 5.02 | -9, 53, -1 |
| 0.096 | 60 | 0.56 | 4.74 | 9, 2, -13 |
|  |  | 0.92 | 4.16 | 15, 8, -16 |
|  |  | 0.99 | 3.62 | 24, 8, -19 |

**Table 3.** Uncertainty processing. Linear relationship (parametric modulation) of activation in the statement phase with uncertainty over all conditions (‘uncertainty’) and in the self-referential condition (‘self uncertainty’). Significance threshold p=0.05 cluster-level FWE-corrected with a cluster-defining threshold (CDT) of p=0.001 unc.

| cluster-level |  | peak-level |  |  |
| --- | --- | --- | --- | --- |
| p<sub>FWE</sub> | k<sub>E</sub> | p<sub>FWE</sub> | t | MNI<br>(mm; x, y, z) |
| **uncertainty** |  |  |  |  |
| <0.001 | 184 | 0.25 | 5.39 | -6, 14, 62 |
|  |  | 0.89 | 4.35 | -9, 23, 38 |
|  |  | 0.99 | 3.76 | 9, 17, 50 |
| 0.036 | 67 | 0.52 | 4.92 | -39, 8, 50 |
| **self uncertainty** |  |  |  |  |
| <0.001 | 155 | 0.095 | 5.93 | -9, 8, 65 |
|  |  | 0.75 | 4.63 | -9, 23, 38 |
|  |  | 0.99 | 3.92 | 9, 23, 38 |
| 0.019 | 76 | 0.23 | 5.46 | -39, 35, -1 |
|  |  | 0.52 | 4.94 | -51, 41, -10 |
|  |  | 0.99 | 4.01 | -48, 50, -10 |
| 0.035 | 65 | 0.34 | 5.23 | 18, 17, 68 |
|  |  | 0.96 | 4.21 | 9, 20, 62 |

**Figure 3.**
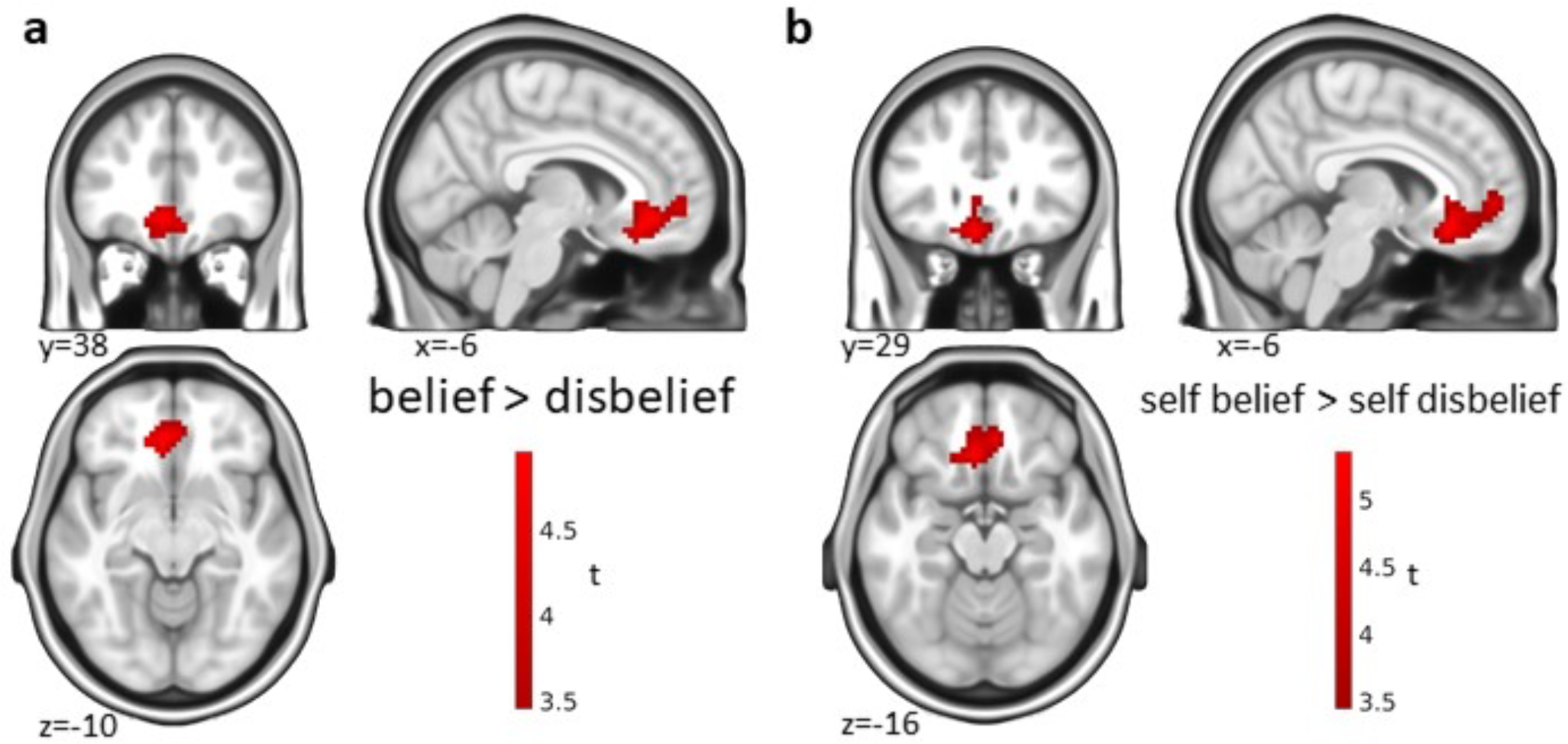
Belief processing. Activation for the contrast ‘belief > disbelief’ in the statement phase a) over all conditions and b) in the self-referential condition. Significance threshold p=0.05 cluster-level FWE-corrected with a cluster-defining threshold (CDT) of p=0.001 unc.

### fMRI uncertainty

We identified a cluster in the dmPFC for uncertainty across all conditions (Figure 4a). Remarkably, our cluster overlapped with the activation identified for uncertainty in Gerchen et al. (2023) (Figure 5b). Again, a within-condition uncertainty effect could only be found in the self condition (Figure 4b) but not for the close or public persons. The cluster for uncertainty in self-referential belief evaluations looked different from the cluster for uncertainty across all conditions in shape but was also localized in the dmPFC. Findings for the opposite contrast direction (certainty) that were also present are shown in the supplement (Supplementary, Figure 2).

**Figure 4.**
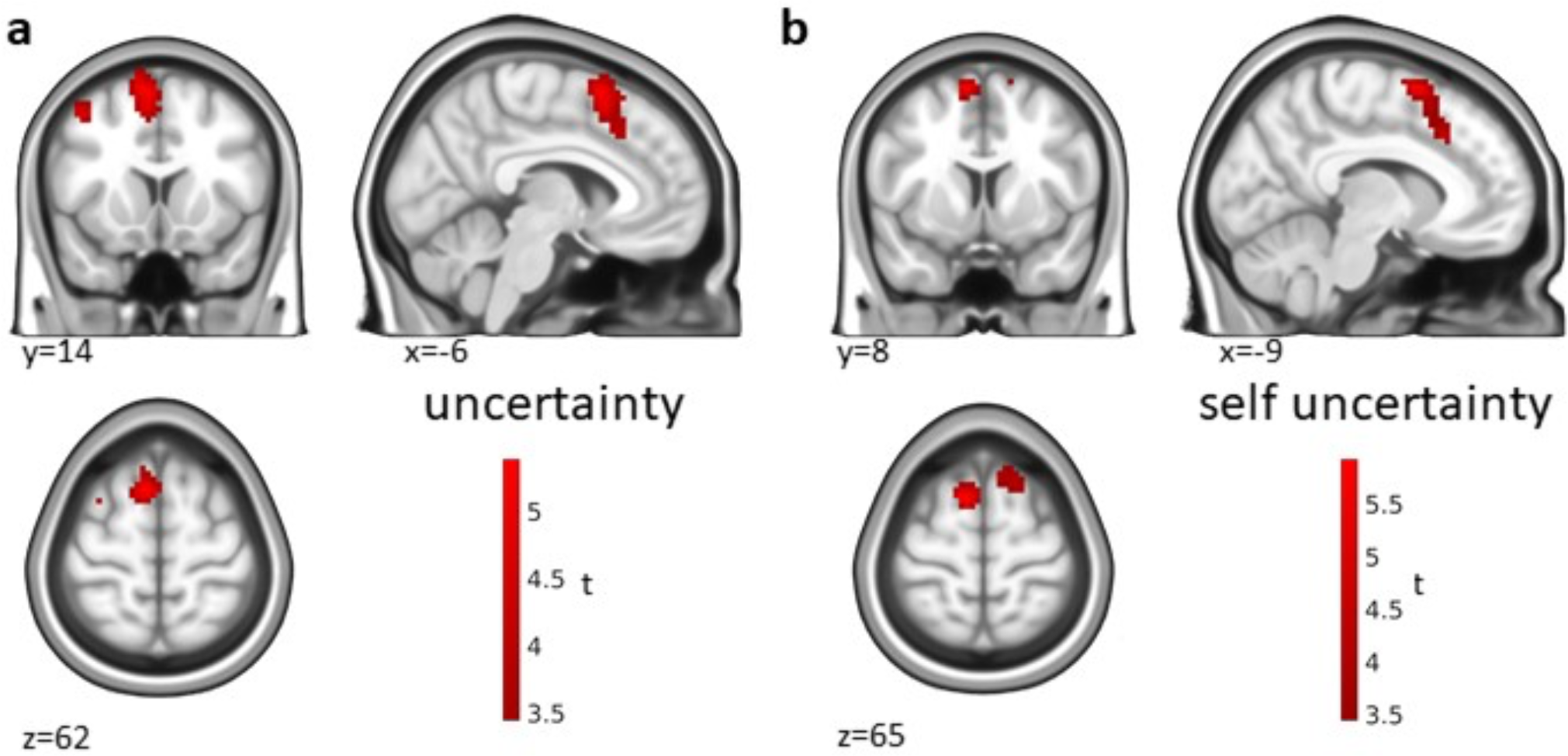
Uncertainty processing. Linear relationship (parametric modulation) of activation in the statement phase with uncertainty a) over all conditions and b) in the self-referential condition. Significance threshold p=0.05 cluster-level FWE-corrected with a cluster-defining threshold (CDT) of p=0.001 unc.

**Figure 5.**
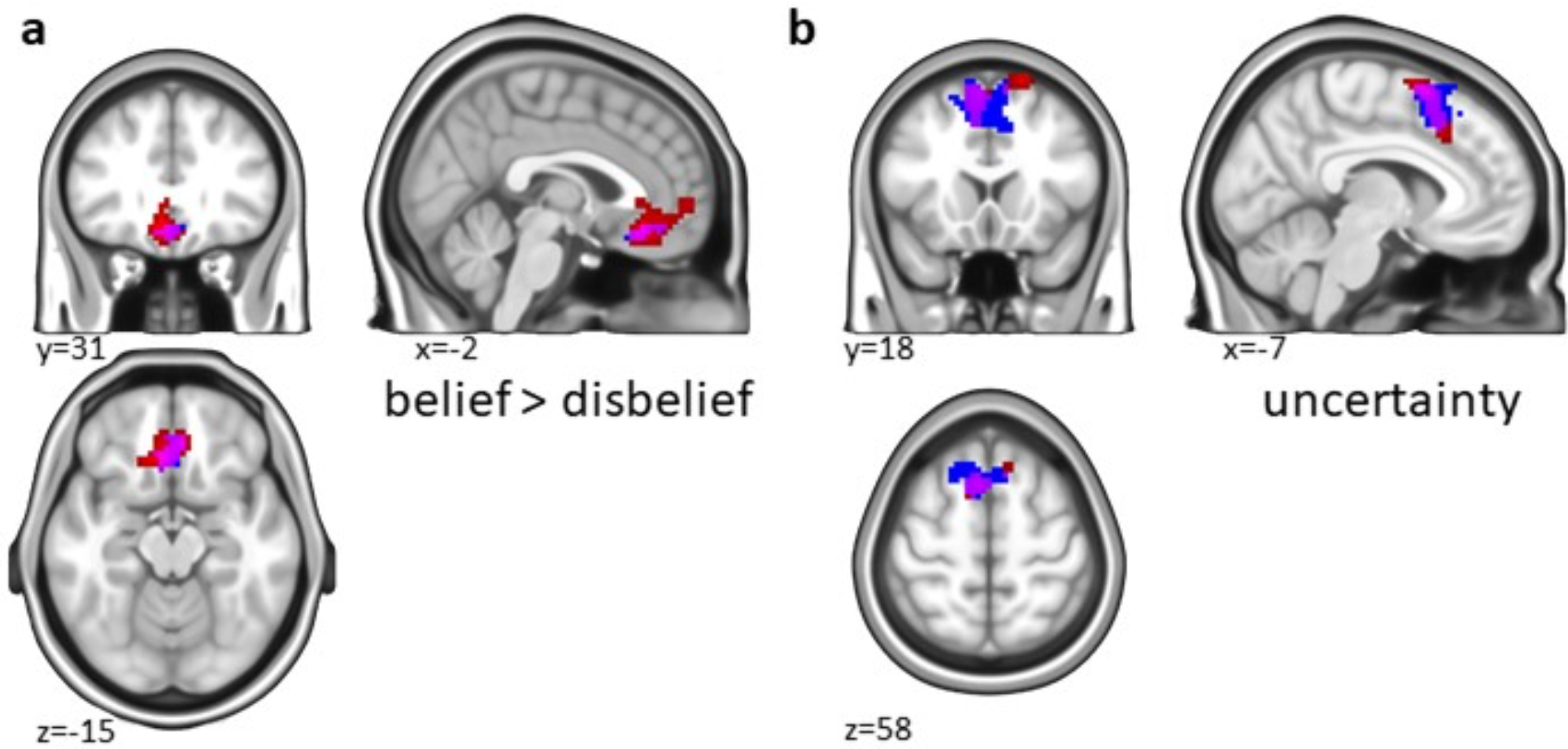
Overlap of self-referential belief processing with general belief processing in Gerchen et al., 2023. a) For the contrast ‘belief > disbelief’. b) For the association with uncertainty. Red: Self-referential belief processing; Blue: General belief processing for statements about facts, politics, religion, conspiracy theories, and superstition reported in Gerchen et al., 2023. Magenta: Overlap of the effects

## Discussion

In the current study, we investigated the question whether the neural correlates of self-referential belief processing differ from those of general belief. For this purpose, we had participants evaluate their belief in statements about themselves, a close, and a public person. While we found belief effects in the vmPFC (Figure 3a) and uncertainty effects in the dmPFC over all conditions (Figure 4a) and within the self-referential condition (Figure 4b), we did not find differences between the conditions, and the effects identified in the self-referential condition were overlapping with our previous effects for general belief processing (Figure 5; Gerchen et al., 2023).

With respect to behavior, in our study the distribution of true versus false statements was slightly skewed across conditions, indicating marginally unbalanced item endorsement. The distribution of statements rated as true versus false also varied between the conditions, with the highest rate of endorsement in the self condition and the lowest in the public person condition. This may be explained by item selection as well as choice of the public person, i.e. the previous German chancellor, who is a Conservative politician while most student participants (and their close peers) might be more progressive.

The certainty ratings also differed between conditions. On average, participants were most certain in their belief evaluations regarding the self-chosen close person and least certain for the public person. The finding that participants placed greater certainty in self-referential judgments than in the judgments of a person who is not a friend or family member is in line with an early report by Kuiper and Rogers (1979). The relatively low certainty in truth evaluations of the public person might be explained by the lack of personal connection and shared episodic memories.

Notably, certainty in belief evaluation was significantly greater than certainty in disbelief evaluations. Interestingly, this finding is inconsistent with the results of Gerchen et al. (2023), where the effect was inverse, which might be due to the different nature of statements used. It might be the case that participants more vehemently disagreed with superstitious or conspiracy statements used in Gerchen et al. (2023), while they were more uncertain in abstract evaluations of themselves or the close and other persons.

As expected, self-referential processing (self > other) revealed a large cluster in the vmPFC and ACC, also reaching into the dmPFC, corpus callosum and subcortical structures (Figure 2).

Belief vs. disbelief evaluation was associated with activation in the vmPFC and ACC (Figure 3a). Within conditions, the contrast ‘belief > disbelief’ was only significant in the self-condition (Figure 3b), with the cluster looking almost identical to the one found over all conditions in location, size, and shape. The lack of significance in the close and public person conditions in our study might be due to low power and stronger neural activation for the self, as target words describing the self rather than another person are salient and self-relevant stimuli (Sui & Humphreys, 2015). This would also be consistent with Gao et al. (2022), who found significantly greater vmPFC activation for self-belief than belief in statements about a gender-matched celebrity. It is, however, important to note that in our case we did not identify a significant difference between the conditions.

Interestingly, the cluster identified for self-referential belief was in close vicinity to the one identified for self-referential processing (self > other) but differed in exact location, size, and shape (Supplementary Figure 3). These distinguishable clusters imply that self-referential processing and belief processing are *distinct* neural processes embedded in the same neural networks, bearing important implications for later investigations in clinical populations displaying aberrant belief and/or self-referential processing.

Importantly, the cluster we identified for self-referential belief replicates the vmPFC belief processing effect in Gerchen et al. (2023; Figure 5a, Supplementary Figure 1). Notably, Gerchen et al. (2023) identified the effect during the decision phase and linked it to decision making, whereas here it was identified during the statement presentation phase. This difference might be related to the higher temporal precision and the simpler statement structure (3 words in comparison to complex statements) in the present study, which could lead to faster and more precise decision processes. Nonetheless, future research is needed to clarify the exact role of the vmPFC in belief processing. In the context of believing and deciding, the vmPFC appears to play a crucial role in belief processing of statements about the self, others, and a variety of other categories (e.g., Harris et al., 2008; Gao et al., 2022; Gerchen et al., 2023), “false tagging” dubious cognitive representations (Asp et al., 2012), signaling emotional saliency (Goel & Dolan, 2003) as well as recruiting somatic markers for decision-making (Bechara et al., 2000; Damasio, 1996).

In sum, our findings suggest a *common neural correlate* for self-referential belief and general belief processing in the vmPFC. As Gao et al. (2022) and our descriptive results show, the vmPFC effect might be even slightly stronger in self-referential processing, but the localization of the effect is clearly the same.

In contrast to Gerchen et al. (2023), we did not identify the left lPFC, left superior parietal cortex or dlPFC for belief processing during statement presentation. Gerchen et al. (2023) suggested that the lPFC’s role in relational integration of working memory contents might lead to its proposed role in generating a belief signal. A reason why we did not find the dlPFC or superior parietal cortex in this study might be the lower working memory load (role of dlPFC in working memory: e.g., Barbey et al., 2013; role of superior parietal cortex in working memory: e.g., Koenigs et al., 2009), as participants only had to retain the target word and person to be evaluated, whereas Gerchen et al. (2023) used more complex statements. On the other hand, the lPFC has been shown to be involved in autobiographical memory search (Cabeza & St Jacques, 2007). It remains thus unclear why it was not identified in this study.

We also did not identify any region for opposite contrast ‘disbelief > belief’, while Gerchen et al. (2023) found the anterior TP. This might be a result of different experimental designs and low power and should be further investigated in future studies. A plausible explanation as to why we did not replicate the anterior TP might be because, contrary to Gerchen et al. (2023), we did not include factual statements of semantic categories and the TP has repeatedly been implicated in semantic processing and memory (Bonner & Price, 2013; Chen et al., 2017; Patterson et al., 2007; Simmons & Martin, 2009; Tsapkini et al., 2011; Vandenberghe et al., 2002).

With respect to the uncertainty/certainty factor of belief postulated by Gerchen et al. (2023), we identified a cluster in the dmPFC associated with uncertainty that was again present over all conditions (Figure 4a) and within the self-referential condition (Figure 4b). Notably, this effect replicates the findings of Gerchen et al. (2023) we interpreted as a “subjective epistemic risk” marker despite the different nature of statements, again implying a *common neural correlate* of self-referential and general belief for this process.

In another context, Kaplan et al. (2016) found that dmPFC activation during challenges to political positions negatively correlated with belief change, which is coherent with its supposed role in uncertainty. This process might thus become relevant in future studies in clinical populations with heightened risk processing and uncertainty (e.g., eating disorders, Brown et al., 2017; anxiety, depressive and obsessive-compulsive disorders; Gentes & Ruscio, 2011), where dmPFC activation might serve as a diagnostic marker or indicator of change processes.

Notably, several limitations apply to our study. First, the sample mainly consisted of university students and is thus not representative of the general population. Second, the analyses were likely underpowered, as we only attained a final sample size of 27 participants. This might play a part in why we did not find significant effects for the contrast ‘belief > disbelief’ within the close and public person conditions, as activation in these regions might be weaker for other-than for self-reference as discussed above. Third, participants might have been less vigilant and conscientious at the time of measurement in the fMRI due to the previous run of the MEG part of the experiment. Fourth, the two parallel sets of target items were not perfectly balanced for the public person statements, which might be due to the limited sample of N=9 for the pre-test. Finally, the distribution of true vs. false and certainty ratings was skewed between the three conditions. Thus, further future well-powered and confirmatory analyses are warranted to corroborate our findings.

## Conclusion

By demonstrating common neural correlates of self-referential and general belief processes in the vmPFC and dmPFC and distinguishing self-referential processing from self-referential belief processing, our findings lay a foundation stone for future studies on belief processing. Further investigating these processes might be especially relevant in clinical populations with aberrant (self-referential) belief (e.g., depression, psychosis) and certainty processing (e.g., anxiety, depressive, eating, and obsessive-compulsive disorders), where they might serve as a diagnostic marker and indicator of intervention related change processes.

## Acknowledgements

GK and MFG were supported by the WIN program of the Heidelberg Academy of Sciences and Humanities financed by the Ministry of Science, Research, and the Arts of the State of Baden-Württemberg. The funding sources were not involved in study design, the collection, analysis and interpretation of data, and in the writing of the manuscript.

## Author contributions

EB, IS, PK, & MFG planned and designed the study. EB, IS, & MFG created the study material. EB, IS, & MFG conducted the study and analyzed the data. EB, GK, PK, & MFG wrote the manuscript.

