## Supplementary_Information for "Neural Correlates of Self-Referential Belief Processes"

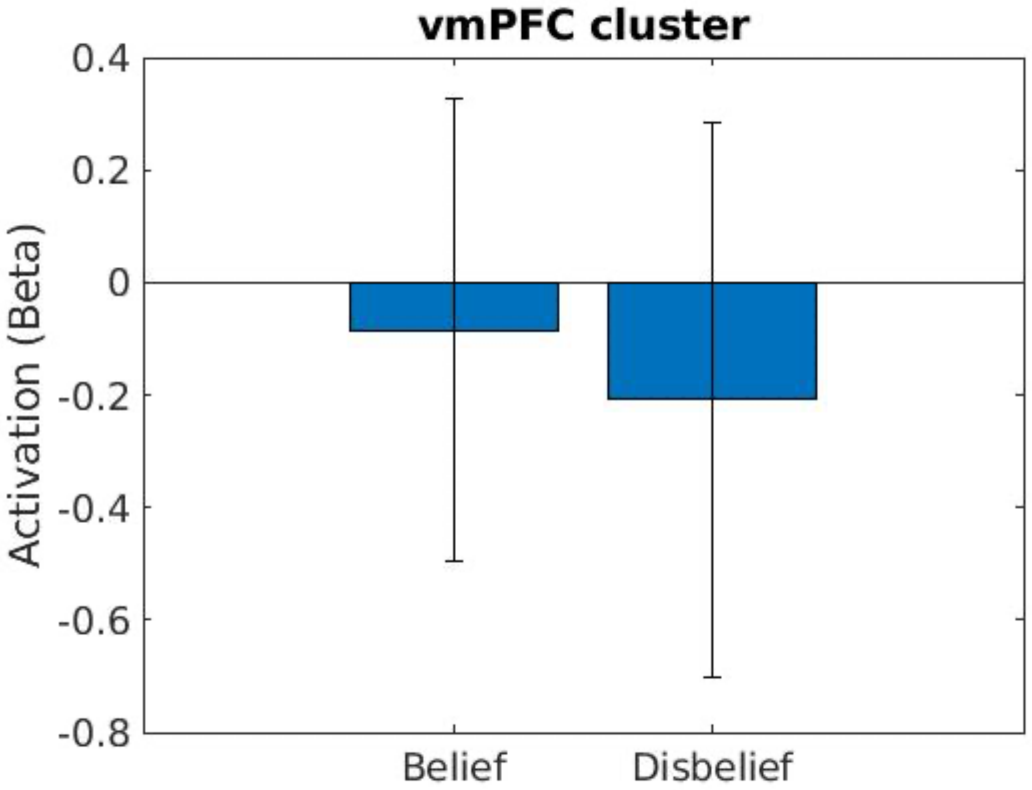


**Supplementary Figure 1. vmPFC Activation.** Condition-specific activation (averaged beta values +/- 1 SD) in the vmPFC cluster in the statement phase (Figure 3a). The positive contrast effect ('belief > disbelief')' is created by a stronger deactivation in the disbelief condition than in the belief condition.

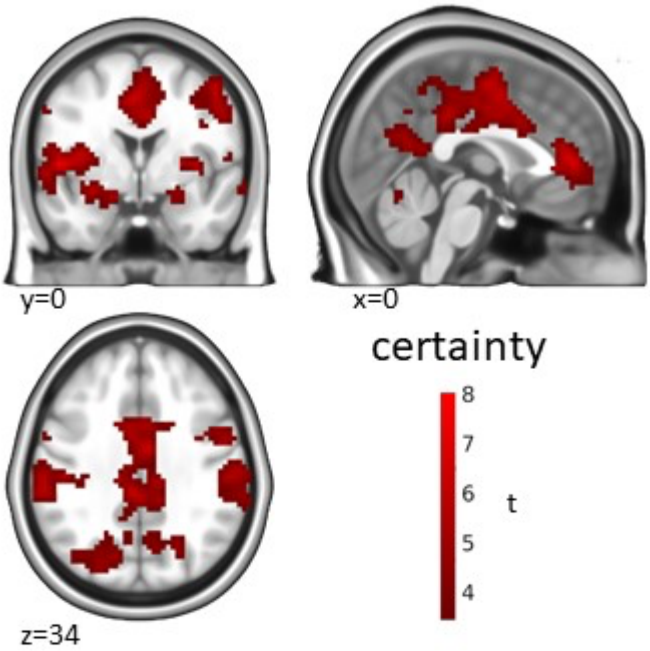


**Supplementary Figure 2. Certainty processing.** Linear relationship (parametric modulation) of activation in the statement phase with certainty over all conditions. Significance threshold p=0.05 cluster-level FWE-corrected with a cluster-defining threshold (CDT) of p=0.001 unc.

| cluster-level | | peak-level | | |
|---|---|---|---|---|
| $p_{FWE}$ | $k_E$ | $p_{FWE}$ | t | MNI (mm; x, y, z) |
| **certainty** | | | | |
| <0.001 | 9845 | <0.001 | 8.03 | -9, -7, 53 |
| | | 0.002 | 7.69 | -48, -4, 5 |
| | | 0.002 | 7.65 | -60, -31, 20 |
| <0.001 | 626 | 0.003 | 7.41 | 3, 44, 8 |
| | | 0.11 | 5.85 | -9, 35, -1 |
| | | 0.13 | 5.73 | -9, 50, -1 |
| 0.002 | 129 | 0.065 | 6.10 | 30, -46, 65 |
| | | 0.77 | 4.58 | 30, -43, 44 |
| | | 0.83 | 4.49 | 24, -43, 53 |
| 0.036 | 67 | 0.30 | 5.29 | 24, -4, -16 |
| | | 0.74 | 4.62 | 12, 2, -13 |
| 0.002 | 122 | 0.52 | 4.92 | 66, -19, -7 |
| | | 0.91 | 4.32 | 63, -4, -10 |

**Supplementary Table 1. Certainty processing.** Linear relationship (parametric modulation) of activation in the statement phase with certainty over all conditions. Significance threshold p=0.05 cluster-level FWE-corrected with a cluster-defining threshold (CDT) of p=0.001 unc.

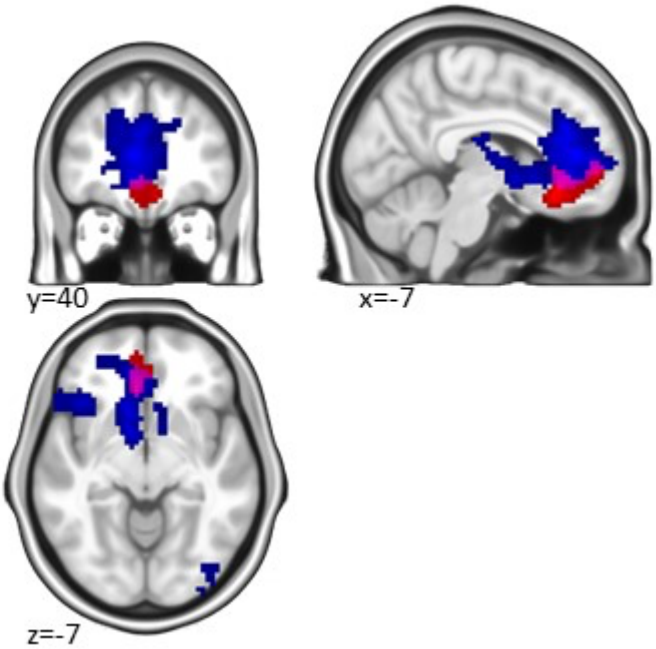


**Supplementary Figure 3. Spatial relationship of activations for self-referential processing and self-referential belief processing.** Blue: Activation for the contrast self > other (Fig. 2.). Red: Activation for the contrast self belief > self disbelief (Fig. 3b). Magenta: Overlap of the effects. While the activated clusters are contiguous and partly overlap at the border regions, under consideration of the effect of intraindividual variation and smoothing on group-level fMRI results they appear to be based on distinct neural substrates. Significance threshold p=0.05 cluster-level FWE-corrected with a cluster-defining threshold (CDT) of p=0.001 unc.

| Item Nr. | Original Item | English Translation | Set |
|---|---|---|---|
| 1 | anpassungsfähig | adaptable | 2 |
| 2 | großzügig | generous | 1 |
| 3 | vergesslich | forgetful | 2 |
| 4 | nachdenklich | pensive | 1 |
| 5 | autoritär | authoritarian | 1 |
| 6 | intelligent | intelligent | 1 |
| 7 | leichtgläubig | gullible | 2 |
| 8 | besorgt | worried | 2 |
| 9 | empfindsam | sensitive | 2 |
| 10 | zuversichtlich | confident | 1 |
| 11 | naiv | naïve | 1 |
| 12 | skeptisch | sceptical | 1 |
| 13 | emotional | emotional | 1 |
| 14 | ehrlich | honest | 2 |
| 15 | rücksichtslos | inconsiderate | 1 |
| 16 | romantisch | romantic | 1 |
| 17 | attraktiv | attractive | 1 |
| 18 | entspannt | relaxed | 2 |

| | | | |
|---|---|---|---|
| 19 | wehleidig | whining | 1 |
| 20 | stur | stubborn | 1 |
| 21 | widerstandsfähig | robust | 2 |
| 22 | gesellig | gregarious | 1 |
| 23 | einsam | lonely | 1 |
| 24 | unberechenbar | unpredictable | 2 |
| 25 | streng | strict | 2 |
| 26 | stark | strong | 1 |
| 27 | liebenswert | kind | 2 |
| 28 | launisch | capricious | 1 |
| 29 | sarkastisch | sarcastic | 2 |
| 30 | dominant | dominant | 1 |
| 31 | taktvoll | tactful | 2 |
| 32 | kompetent | competent | 1 |
| 33 | unzufrieden | dissatisfied | 2 |
| 34 | gehemmt | inhibited | 1 |
| 35 | rational | rational | 1 |
| 36 | tolerant | tolerant | 2 |
| 37 | pessimistisch | pessimistic | 1 |
| 38 | schreckhaft | jumpy | 2 |

| | | | |
|---|---|---|---|
| 39 | bescheiden | modest | 1 |
| 40 | faul | lazy | 1 |
| 41 | normal | normal | 1 |
| 42 | provokant | provocative | 1 |
| 43 | selbstsicher | self-confident | 2 |
| 44 | loyal | loyal | 2 |
| 45 | impulsiv | impulsive | 2 |
| 46 | beharrlich | persistent | 2 |
| 47 | konsequent | consistent | 2 |
| 48 | ruhelos | restless | 1 |
| 49 | idealistisch | idealistic | 1 |
| 50 | verständnisvoll | understanding | 2 |
| 51 | liebevoll | loving | 2 |
| 52 | langweilig | boring | 1 |
| 53 | sparsam | thrifty | 2 |
| 54 | humorvoll | humorous | 2 |
| 55 | nachlässig | negligent | 2 |
| 56 | verträumt | dreamy | 1 |
| 57 | interessant | interesting | 1 |
| 58 | sprunghaft | erratic | 2 |

| | | | |
|---|---|---|---|
| 59 | sanft | gentle | 1 |
| 60 | unterhaltsam | entertaining | 2 |
| 61 | mutig | courageous | 1 |
| 62 | unflexibel | inflexible | 2 |
| 63 | unkonventionell | unconventional | 2 |
| 64 | unkompliziert | uncomplicated | 1 |
| 65 | ungeduldig | impatient | 1 |
| 66 | abenteuerlustig | adventurous | 2 |
| 67 | beliebt | popular | 1 |
| 68 | unpünktlich | unpunctual | 2 |
| 69 | stolz | proud | 2 |
| 70 | wissbegierig | inquisitive | 1 |
| 71 | berechnend | calculating | 2 |
| 72 | intellektuell | intellectual | 2 |
| 73 | gesprächig | talkative | 1 |
| 74 | ordentlich | tidy | 2 |
| 75 | ängstlich | anxious | 2 |
| 76 | ehrgeizig | ambitious | 2 |
| 77 | fleißig | hard-working | 1 |
| 78 | stabil | stable | 2 |

| | | | |
|---|---|---|---|
| 79 | unsicher | uncertain | 1 |
| 80 | materialistisch | materialistic | 2 |

**Supplementary Table 2. Study materials.** List of adjectives used in the task in original German and translated to English. 'Set' refers to the adjective set the word was assigned to.

| | Set 1 Mean ± StD | Set 2 Mean ± StD | t | DoF | p | Hedges' g |
|---|---|---|---|---|---|---|
| **self-referential belief ratio** | 0.68 ± 0.07 | 0.63 ± 0.11 | 1.45 | 25 | 0.16 | 0.54 |
| **self-referential certainty** | 76.44 ± 6.53 | 78.45 ± 10.17 | -0.60 | 25 | 0.55 | -0.23 |
| **close person belief ratio** | 0.65 ± 0.08 | 0.62 ± 0.10 | 0.80 | 25 | 0.43 | 0.30 |
| **close person certainty** | 76.97 ± 6.36 | 81.59 ± 10.40 | -1.38 | 25 | 0.18 | -0.52 |
| **public person belief ratio** | 0.58 ± 0.07 | 0.49 ± 0.05 | 3.85 | 25 | <0.001 | 1.44 |
| **public person certainty** | 58.47 ± 4.93 | 65.54 ± 14.11 | -1.26 | 25 | 0.22 | -0.47 |

**Supplementary Table 3. Comparison of adjective sets.** Statistical comparison of responses (belief ratio: ratio of 'yes' to 'no' answers and certainty) in the fMRI experiment between the two adjective sets that were used in the parallel versions of the task.
